# Dedicated nanoparticle flow cytometry for single extracellular vesicle phenotyping: Performance of the CytoFLEX Nano

**DOI:** 10.64898/2025.12.15.694348

**Authors:** Johann Mar Gudbergsson, Mette Galsgaard Malle, Josefine Bech Jensen, Elisabeth Noble Truelsen, Jonas Klejs Hemmingsen, Samanta Nelson, Jørgen Kjems, Aida Solhøj Hansen, Anders Etzerodt, Anja Bille Bohn

**Affiliations:** Department of Biomedicine, Aarhus University, Aarhus, Denmark; Interdisciplinary Nanoscience Center, Aarhus University, Aarhus, Denmark; Department of Molecular Medicine, University of Southern Denmark, Odense, Denmark; Department of Clinical Medicine, Aarhus University, Aarhus, Denmark; Department of Infectious Diseases, Aarhus University Hospital, Aarhus, Denmark; Department of Molecular Biology and Genetics (MBG), Aarhus University, Aarhus, Denmark; Department of Rheumatology, Aarhus University Hospital, Aarhus, Denmark

**Keywords:** nanoflow, exosome, microvesicle, high-throughput, staining

## Abstract

Accurate discrimination of extracellular vesicles (EVs) from non-vesicular nanoparticles and robust phenotyping of individual EVs directly within complex biofluids are essential to advance understanding of EV biology and to realize their potential as biomarkers and therapeutic agents. Conventional flow cytometers, originally designed for cellular analysis, lack the scatter and fluorescence sensitivity, dynamic range, and event-rate control required for quantitative characterization of nanoscale vesicles and are particularly susceptible to coincident (swarm) detection. Dedicated nanoparticle flow cytometers have been developed to address these limitations, and here we systematically evaluate the suitability of the CytoFLEX Nano (Beckman Coulter) for EV analysis. Using a series of calibration beads of different materials, fluorescent liposomes, and EVs isolated from peritoneal fluid and cell culture medium, we assess size-detection thresholds, scatter and fluorescence sensitivity, dynamic range, and volumetric counting accuracy. These data provide practical guidance for implementing nanoscale flow cytometry in EV research and support the informed adoption of dedicated nanoparticle cytometers in studies adhering to current EV reporting standards.

## Introduction

Extracellular vesicles (EVs) comprise a highly heterogeneous population of lipid bilayer–enclosed particles, ranging in size from approximately 30□nm to 10□µm in diameter. They are released by virtually all cell types and found in all major biological fluids under both physiological and pathological states. Over recent years, small EVs (sEVs,□<□200□nm) have emerged as pivotal mediators of intercellular communication, modulating immunity, tissue regeneration, and homeostasis, and are implicated in the progression of disorders such as cancer, neurodegeneration, and chronic inflammatory diseases [1], [2], [3]. Their ability to transfer proteins, nucleic acids, lipids, and metabolites to recipient cells underscores their potential as disease biomarkers and therapeutic targets [4], [5]. sEVs originate from either direct plasma membrane budding (microvesicles) or via endosomal pathways (exosomes), including late endosomes and autophagy-related compartments, carrying molecular cargo that reflects their cellular origin and biogenic route [6], [7], [8]. The inherent heterogeneity in the size, density, composition, and biogenesis of sEVs complicates their accurate classification, and the co-isolation of non-vesicular particles such as lipoprotein particles and protein aggregates remains a significant source of analytical ‘noise’ [9], [10], [11].

Among available analytical techniques, flow cytometry stands out for its ability to perform rapid, high-throughput, multiparametric analysis of EVs at the single-particle level. Compared to nanoparticle tracking analysis (NTA) and electron microscopy (EM), flow cytometry allows simultaneous physical and phenotypic characterization of thousands of sEVs, making it a promising tool for subtype profiling. However, conventional flow cytometers face substantial challenges; the small size and low refractive index of sEVs require highly sensitive optics, optimized instrument settings, extensive cleaning programs, and robust controls to distinguish true vesicular events from background or co-precipitated particles. Even with these measures, size limits for the most sensitive conventional flow cytometers typically extend to approximately 80–120□nm for polystyrene (PS) beads and even higher for sEVs given their lower refractive index [12], [13]. Often fluorescence triggering is used to improve sensitivity to detect EVs that are below the size (scatter) threshold. However, a general methodological concern for EV flow cytometry is swarm detection of multiple EVs as single events, making rigorous dilution controls essential.

To address these challenges, dedicated nanoparticle flow cytometers have been developed to improve the detection and characterization of sEVs. In early□2024, Beckman□Coulter launched the CytoFLEX□Nano, advertising particle detection down to 40□nm PS beads. In this study, we systematically evaluate the performance of the CytoFLEX□Nano in terms of scatter, fluorescence resolution and sensitivity, by analyzing a panel of reference beads and biological samples. Our aim is to highlight the potential of the instrument as well as share challenges observed during our testing of this high-resolution and high-throughput instrument

## Materials & methods

### Nanoparticle flow cytometry

Samples were acquired on the CytoFLEX Nano, which is equipped with four lasers (405 nm, 488 nm, 561 nm, and 638 nm) and 6 fluorescence detectors, 1 forward scatter detector and 5 side scatter detectors (Beckman Coulter, Souzhou, China) using CytExpert nano software (v. 1.1.0.6, Beckman Coulter, Souzhou, China). Before each experiment, the daily QC, including control of both scatter parameters and fluorescence parameters, was acquired to ensure optimal instrument performance. After adjusting the gain settings for each experiment, a buffer sample was acquired and used to adjust threshold to an event rate of 20-200 events/sec (threshold varied between 300 and 450 depending on buffer and experiment). Beads were diluted in freshly opened sterile PBS and optimal dilutions were found before data acquisition. The list and dilution of beads used here can be found in Table 1.

**Table 1.** Overview of beads used in this study.

| Catalogue number | Manufacturer | Material | Description | Bead diameter (nm) | Dilution |
| --- | --- | --- | --- | --- | --- |
| 20812125 | PolyAn | PMMA | SCB250416AM4 – SCB (4 peaks) - Carboxy | 100 - 150 | x 10.000 |
| 20812225 | PolyAn | PMMA | SCB250416BM4 – SCB (4 peaks) - Carboxy | 201 - 250 | x 1000 |
| 20812500 | PolyAn | PMMA | SCB250416CM4 – SCB (4 peaks) - Carboxy | 475 - 525 | x 1000 |
| - | PolyAn | PMMA | SCB 0 (Single fluorescence peak) - blank | 120 | x 10.000 |
| - | PolyAn | PMMA | SCB 10 (Single fluorescence peak) | 120 | x 10.000 |
| - | PolyAn | PMMA | SCB 50 (Single fluorescence peak) | 120 | x 10.000 |
| - | PolyAn | PMMA | SCB 500 (Single fluorescence peak) | 150 | x 10.000 |
| - | PolyAn | PMMA | SCB 0 (Single fluorescence peak) - blank | 220 | x 1000 |
| - | PolyAn | PMMA | SCB 10 (Single fluorescence peak) | 240 | x 1000 |
| - | PolyAn | PMMA | SCB 50 (Single fluorescence peak) | 230 | x 1000 |
| - | PolyAn | PMMA | SCB 0 (Single fluorescence peak) - blank | 470 | x 1000 |
| - | PolyAn | PMMA | SCB 10 (Single fluorescence peak) | 490 | x 1000 |
| - | PolyAn | PMMA | SCB 50 (Single fluorescence peak) | 500 | x 1000 |
| NFPPS-0152-5 | Spherotech | PS | Single fluorescence peak | 130 | x 10 |
| RCP-05-5 | Spherotech | PS | 4 fluorescence peaks (4-peak) | 400 - 600 | x 10 |
| D03231 | Beckman Coulter | PS | NanoVIS Low Scatter calibration beads (4 scatter peaks) | 40, 80, 108, 142 | x 1 |
| D03231 | Beckman Coulter | PS | NanoVIS High Scatter calibration beads (4 scatter peaks) | 142, 304, 600, 1020 | x 1 |
| 8050 | Thermo Scientific | Silica | 8000 Series Silica Particle Size Standards | 500 | x 100.000 |
| 8070 | Thermo Scientific | Silica | 8000 Series Silica Particle Size Standards | 700 | x 100.000 |
| 8100 | Thermo Scientific | Silica | 8000 Series Silica Particle Size Standards | 1000 | x 100.000 |

Data from all samples and controls within an experiment was acquired at the same speed and stopping conditions to enable comparison between samples. For all EV experiments, several controls were used, including PBS, PBS + stain, single stains, unstained samples, and detergent-treated stained samples. A compilation of controls in EV experiments is represented in Figure S1A-D. Further details on the EV experiments are available in the respective sections below.

### Scatter calibration using FCMPASS

Scatter calibration of data collected with the CytoFLEX Nano was performed using NanoVis polystyrene reference beads (Beckman Coulter) analysed with FCMPASS software to relate light scattering intensity to particle size via Mie theory, as described elsewhere [14], [15]. Initially, NanoVis High beads (nominal diameters: 142 nm, 304 nm, 600 nm, and 1020 nm) were acquired to determine the median signal intensities of each peak in the violet Side Scatter 2 (vSSC2) channel. These values were used to calculate the optical collection half-angle of the instrument, ensuring a fit confidence greater than 90%. This empirically derived half-angle was subsequently applied to calibrate the vSSC1 detector using NanoVis Low beads (nominal diameters: 40 nm, 80 nm, 108 nm, and 142 nm), thereby establishing a standardized relationship between scattering signal and particle diameter across the relevant detection range. Plots from the FCMPASS report on the model fit are shown in Figure S1E.

### Preparation of Atto-conjugated liposomes

Liposomes were prepared from a mixed lipid composition containing 1 % 1,2-dioleoyl-sn-glycero-3-phospho-(1’-rac-glycerol) (DOPG), between 0 to 0.5 % fluorescent lipid (see table below), and remaining 98.5-99% 2-Dioleoyl-sn-glycero-3-phosphocholine (DOPC) given in mole percentage. All Atto dyes are 1,2-Dioleoyl-sn-glycero-3-phosphoethanolamine (DOPE) lipid from Attotec.

All lipids were suspended in chloroform, mixed, evaporated using a N_2_ stream and subsequently dried under vacuum for 1 hour. The lipid film was hydrated in filtered PBS (Dulbecco′s Phosphate Buffered Saline), vortexed and incubated for 30 min at room temperature (RT). The liposomes were frozen and thawed for 10 cycles to ensure unilammelarity and extruded at 50 nm using an Avanti Mini-extruder.

The same mole percentage of Atto-DOPE lipids were used for controls, to show that no micellar formation was observed. The lipids were evaporated using N_2_ and placed in vacuum for 1 hour. The lipid film was hydrolysed in filtered PBS (Dulbecco′s Phosphate Buffered Saline), vortexed, and incubated for 30 min at RT. For the micellar control, lipids were evaporated under N□ and then placed under vacuum for 1 h. The dry lipid film was hydrated in filtered PBS (Dulbecco’s phosphate-buffered saline), vortexed, and incubated for 30 min at RT.

The mole percentage of fluorescent lipids can be converted to an estimated number of lipids from the following equation:

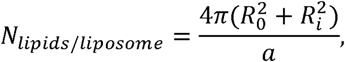

Where R_0_ = outer liposome radius, R_i_ = the inner liposome radius given by R_0_-t, with t being the membrane thickness (approximately 4 nm). The molecular area per lipid, a, is estimated to 0.7 nm^2^ at RT, corresponding primarily to DOPC. Table 3 shows calculated number of lipids per liposome for diameters ranging from 50 nm (extruded diameter) to 70 nm (measured hydrodynamic radius) and the corresponding number of fluorescent lipids for 0.5% and 0.05%.

**Table 2.** Lipid composition and fluorochrome concentrations in liposomes.

| Sample name | Atto655-DOPE | Atto565-DOPE | Atto488-DOPE | DOPG | DOPC |
| --- | --- | --- | --- | --- | --- |
| **15 Liposome-Atto655** | 0.05 % |  |  | 1 % | 98.5 % |
| **30 Liposome-Atto655** | 0.1 % |  |  | 1 % | 98.83% |
| **50 Liposome-Atto655** | 0.167 % |  |  | 1 % | 98.9 % |
| **150 Liposome-Atto655** | 0.5% |  |  | 1 % | 98.95 % |
| **15 Liposome-Atto565** |  | 0.05 % |  | 1 % | 98.5 % |
| **30 Liposome-Atto565** |  | 0.1 % |  | 1 % | 98.83% |
| **50 Liposome-Atto565** |  | 0.167 % |  | 1 % | 98.9 % |
| **150 Liposome-Atto565** |  | 0.5% |  | 1 % | 98.95 % |
| **15 Liposome-Atto488** |  |  | 0.05 % | 1 % | 98.5 % |
| **30 Liposome-Atto488** |  |  | 0.1 % | 1 % | 98.83% |
| **50 Liposome-Atto488** |  |  | 0.167 % | 1 % | 98.9 % |
| **150 Liposome-Atto488** |  |  | 0.5% | 1 % | 98.95 % |
| **0 Blank Liposome** |  |  |  | 1 % | 99 % |

**Table 3.** Overview of membrane dyes applied.

| Liposome Diameter | $N_{lipids/vesicle}$ | # fluorochromes for 0.5% | # fluorochromes for 0.05% |
| --- | --- | --- | --- |
| 50 nm | $1.91 \cdot 10^4$ | 96 | 10 |
| 60 nm | $2.83 \cdot 10^4$ | 141 | 14 |
| 65 nm | $3.35 \cdot 10^4$ | 168 | 17 |
| 70 nm | $3.92 \cdot 10^4$ | 196 | 20 |

The measured hydrodynamic radius at 70 nm is larger than the actual mean size and the extruded 50 nm would be smaller than the actual liposome size. However, liposomes are heterogenous in size and lipid composition follow a log-normal distribution [16], [17]. For subsequent analyses we will refer to the liposome with 0.5% dye as approximately 150 fluorophores and the 0.05% dye as approximately 15 fluorophores despite the inherent broad distribution.

### ImageStream analysis

Samples were acquired using the INSPIRE® software (V200.1.620.0, Cytek Biosciences) on an ImageStream X Mk II (Cytek Biosciences) at a 1:10.000 dilution using a 60x magnification, High Gain mode, a core diameter of 7 µm, with the 785 nm laser adjusted to 70mW, and with the “remove beads function” turned off. Atto488, Atto565, and Atto655 were excited by the 488 nm laser (200 mW), 561 nm laser (200 mW), and 642 nm laser (150 mW) and were collected in channel (Ch) 02, Ch04, and Ch11, respectively. Brightfield (BF) images were acquired in Ch01 and Ch09 and Darkfield/sidescatter (SSC) in Ch06. Each sample was acquired for 2 minutes.

Data were analysed in the IDEAS® software (V6.3.41.0, Cytek Bioscience). Liposomes were initially gated as events with a SSC below 1000. To ensure that median values for Atto-positive liposomes were based on single events, a peak mask covering the pixels with highest intensity in Ch02, Ch04, and Ch11, was created and combined with the spot count feature. This allowed us to select single liposomes positive for an Atto dye but was not applicable to the 0% samples. A gate was set above the peak corresponding to 0% liposomes in an intensity Atto488, intensity Atto565, or intensity Atto655 histogram, respectively and transferred to the stained samples as well as to the following controls; PBS alone, PBS + fluorochrome, and to liposome + fluorochrome + detergent.

### Peritoneal lavage and ultracentrifugation

Peritoneal lavage samples were harvested as previously described [18]. Briefly, eight to ten weeks old C57BL6/Rj mice were euthanized by cervical dislocation and 5 mL of PBS with 2 mM EDTA (Invitrogen, #15575020) was injected i.p. The abdomen was massaged for 5-10 s prior to extraction of the fluid. A small incision was made in the skin and blunt dissection done to expose the outer muscle layer. The lateral skin on the right side of the mouse was left attached to allow for stretching of the abdominal muscle layer and displace the abdominal organs with a finger to create an intraperitoneal space for fluid aspiration. A 25G needle was inserted into the peritoneal cavity rostrally pointing caudally and fluid was slowly aspired, and the samples were kept on ice. The peritoneal fluid was centrifuged at 400 x g for 5 min at 4°C to pellet cells and the supernatants were poured into new tubes and centrifuged at 2000 x g for 15 min at 4°C to pellet cells debris. The supernatants were transferred to a new 15 mL tube and stored at -20°C until further analysis.

For ultracentrifugation, 1.5 mL of peritoneal lavage sample was diluted to 20 mL in freshly opened sterile PBS and transferred to an ultracentrifuge tube. Tubes were loaded into a Fiberlite F50L-8 x 39 (Thermo Scientific) fixed-angle rotor and ultracentrifuged at 130.000 x g for 70 min. at 4°C with deceleration set to 4 in a Sorvall Discovery 100 ultracentrifuge. Supernatants were discarded and pellets were resuspended in 1.5 mL PBS.

### EV staining and nano flow cytometry (peritoneal lavage)

Fifty microliters of sample was added to a 96-well plate and 50 µL of 1 µg/mL anti-CD9-AF647 (#124809, BioLegend) antibody diluted in PBS was added to the samples. Antibody vials were centrifuged at 16,000 x g for 5 min prior to preparation to minimize antibody aggregates. The samples were incubated overnight at 4°C. The next day, EV stain mix containing ExoBrite 410/450 CTB EV Stain (Biotium) and ExoBrite 515/540 True EV Membrane Stain (Biotium) was added in 50 µL PBS to the samples at a working concentration of 0.5-1X and incubated 1-2 hours at RT. Samples were then diluted in freshly opened sterile PBS to a final dilution of 1:200 and run on the CytoFLEX Nano (Beckman Coulter). A vehicle (PBS) control sample was run to set the vSSC1 threshold, which was set at 300 for all experiments. Scatter detector gains were kept according to QC values, but relevant fluorescent detectors were set to a gain of 3000. Single stain and unstained controls were acquired to setup compensation. PBS + stain control was run as a background control and for all quantifications the background values were subtracted using the PBS + stain control. A detergent control was used where 2 % Triton X-100 (Sigma Aldrich) was prepared and sterile filtered, and added 1:1 to an already analysed sample and incubated for 30 min at RT. The sample was then analysed to show that the particles that fall into our EV gate are in fact sensitive to detergents. Sample were acquired for 60s at flow rates between 1 – 3 µL/min. Compensation and data analysis was performed in CytExpert Nano (Beckman Coulter) and FlowJo^TM^ v10.10.0 (BD Life Sciences). Quantitative graphs and statistical analyses were created in GraphPad Prism 10 (Dotmatics).

### Cell culture

All cells were maintained in a humidified incubator at 37℃ with 5% CO_2_. The murine mammary carcinoma cell line EO771, originally isolated from the mammary gland of C57BL/6 mice, was purchased from ATCC and cultured in Dulbecco’s Modified Eagle’s Medium (DMEM; 4.5 g/L glucose; Dominique Dutscher) supplemented with 1% exosome-depleted fetal bovine serum (FBS, Gibco) and 1% Penicillin-Streptomycin (Gibco).

### EV isolation by size exclusion chromatography

A total of 7.5 X 10^5^ cells were seeded per T75 culture flask and incubated for 48 hours. Cell culture-conditioned medium (CCM) from 3 culture flasks were harvested, and cells were briefly washed in sterile PBS (Biowest) and added to the harvested CCM. The CCM was centrifuged at 1000 x g for 10 minutes at 4□ followed by centrifugation at 2000 x g for 20 minutes at 4□ to remove cellular debris and apoptotic bodies. A total of 20 mL precleared CCM or non-conditioned medium, used as a negative control, was concentrated at RT with Amicon Ultra-15 Centrifugal filters (Millipore) with a 10 kDa molecular weight cut-off. The concentrated CCM and control medium (500 µL) were applied to size exclusion chromatography columns (qEVoriginal 35 nm Gen 2; Izon Science) and eluted in PBS. Fractions of 500 µL were collected and fractions 1 – 6, 7 – 10, 11 – 12, and 13 – 25 were subsequently pooled for analysis.

### EV staining and nano flow cytometry (culture medium)

Pooled fractions were stained with 0.5X ExoBrite 515/540 True EV Membrane Stain (Biotium) and 0.5X ExoBrite 410/450 CTB EV Stain (Biotium), or with PBS as a control and incubated in the dark for 2 hours at RT. As controls we used PBS alone, PBS + stain (0.5X ExoBrite 410/450 CTB EV Stain and 0.5X ExoBrite 515/540 True EV Membrane Stain) and non-conditioned media + stain. Prior to analysis the stained fractions were diluted 1:10 in PBS. For detergent lysis, the remaining of the recorded samples were treated with 1% Triton X-100 (Sigma-Aldrich) in PBS and incubated at RT for 30 minutes. Samples were analysed using the CytoFLEX Nano. All samples and controls were acquired over a 4-minute period using a flowrate at 1 µL/min. and a vSSC1-H threshold of 450. Detector gains were set to 3000 for V447, B531, Y595, R670, R710 and R792. The gains for forward and side scatter were as follows: vFSC, 1069; vSSC1, 100; vSSC2, 225; bSSC, 100; ySSC, 100; rSSC, 159.

### Staining with different dyes

For the qualitative assessment of different dyes, peritoneal fluid samples were stained as described above. All samples were run in triplicates, and samples were stained according to Table 4.

**Table 4.** Overview of membrane dyes applied.

| Dye | Catalogue / Manufacturer | Stock conc | Working conc. | Excitation | Emission | Final Dilution |
| --- | --- | --- | --- | --- | --- | --- |
| DiO | #60011, Biotium | 1 mM | 1 uM | 484 | 501 | 250 |
| Memglow 640 | #MG04, Cytoskeleton | 20 uM | 0.1 uM | 650 | 673 | 200 |
| ExoBrite CTB | #30111, Biotium | 500x | 0.5 - 1x | 416 | 452 | 500 |
| ExoBrite True EV | #30129, Biotium | 500x | 0.5 - 1x | 515 | 540 | 500 |

## Results

### Separation of various fluorescence intensities on beads of different sizes

Initially, we sought to validate the capability of the CytoFLEX Nano to resolve small polystyrene (PS) beads, as advertised by Beckman Coulter. The instrument is equipped with two violet side scatter (vSSC) detectors, with vSSC1 offering high sensitivity specifically optimized for detecting nanoparticles below 200 nm in diameter. We evaluated NanoVIS beads, spanning sizes from 40 to 1020 nm, and observed clear separation of all expected size-associated peaks using the two vSSC detectors (Figure 1A). To assess fluorescence resolution for small particles, we analysed PS beads (400–600 nm) with four distinct fluorescence intensity peaks (Spherotech). The CytoFLEX Nano efficiently resolved all four peaks across four separate fluorescence channels (Figure 1B). We next examined polymethyl methacrylate (PMMA) beads produced by PolyAn, available in varying fluorescence intensities (SCB 0 to SCB 500, arbitrary unit) and multiple sizes. When analyzing the 120 nm PMMA beads individually, we detected clear separation of fluorescence peaks in the green (B531-H), yellow (Y595-H), and red (R670-H) channels; however, resolution was insufficient when a 4-peak mixture was analysed (Figure 1C). For the 230 nm beads, the 4-peak mixture showed improved resolution in the red channel but remained suboptimal in the green and yellow channels. By contrast, the 500 nm beads displayed well-resolved peaks in both the green and red channels (Figure 1C).

**Figure 1.**
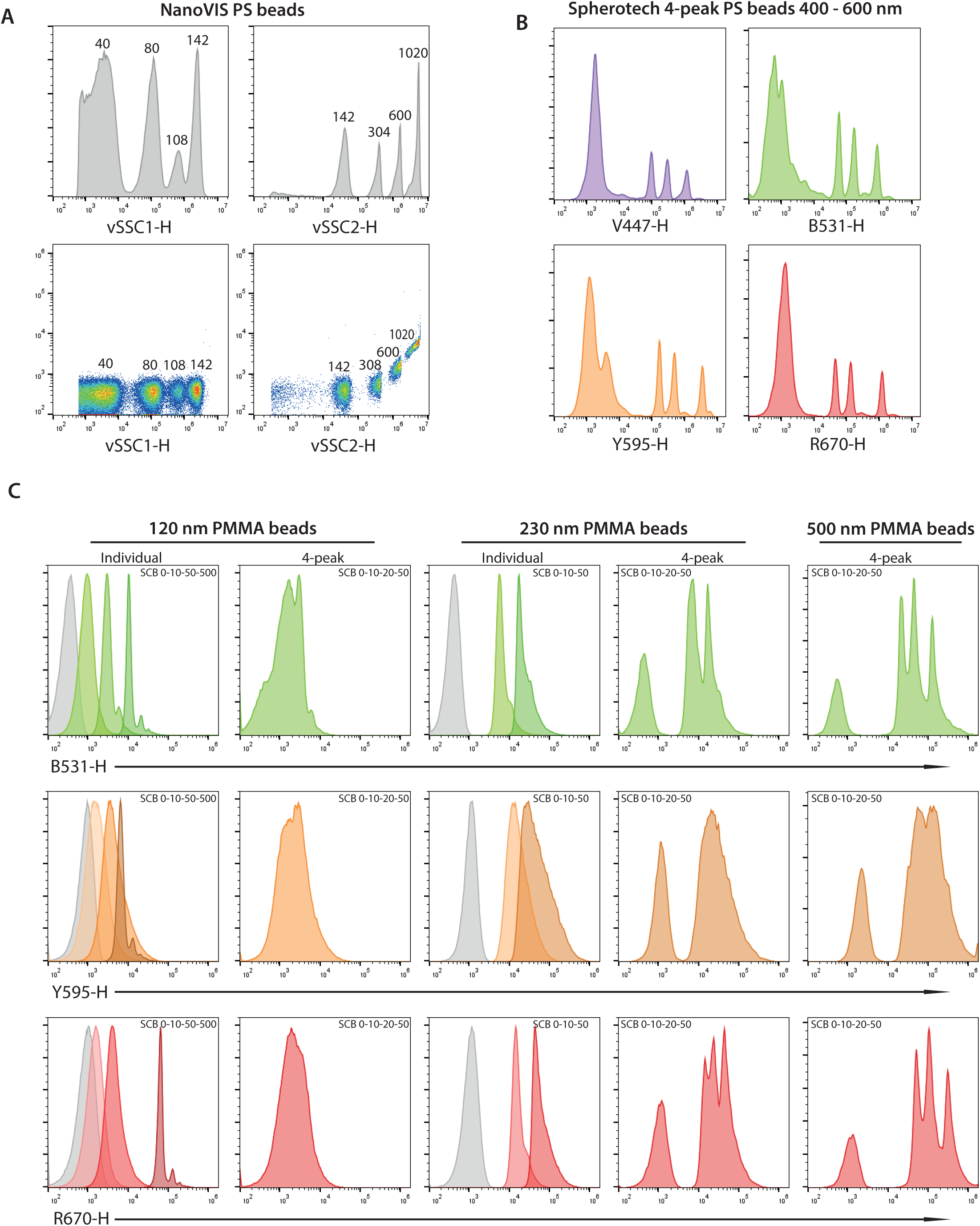
Scatter and fluorescence sensitivity of the CytoFLEX Nano. A) NanoVIS scatter calibration beads shown as histograms and bivariate plots. B) 4-fluorescence peak PS beads from Spherotech, 400 – 600 nm in diameter. C) Individual and gathered 4-peak fluorescent PMMA beads from PolyAn of different sizes.

### Liposomes with differing numbers of fluorochromes as a reference material closer to EVs

Given that the 120 nm PolyAn PMMA beads represented the smallest available 4-peak fluorescence calibration beads and lacked quantitative information regarding their fluorochrome content, we generated synthetic liposomes incorporating varying numbers of fluorophores specific to the green, yellow, and red channels. Liposomes were prepared containing Atto488, Atto595, or Atto655, with an average fluorophore incorporation of 0, 15, 30, 50, or 150 molecules per vesicle as described above (Figure 2A, Tables T1 & T2). Each preparation was analysed on the CytoFLEX Nano in technical triplicates. Fluorescence intensity peaks corresponding to 15, 50, and 150 fluorophores per liposome were clearly resolved, while the population with 30 fluorophores overlapped with those of 15 and 50, impairing distinct peak separation (Figure 2B). Analysis of median fluorescence intensity (MFI) for each preparation revealed a linear correlation, confirming the instrument’s capability for quantitative discrimination of fluorescence intensity among small liposomes (Figure 2C). Upon mixing fluorescently labeled and blank liposomes, populations were effectively distinguished as two discrete peaks corresponding to unstained and stained liposomes (Figure 2D). To exclude the possibility of size heterogeneity influencing fluorescence separation, side scatter calibration was performed to estimate hydrodynamic diameter via FCMPASS software, modeling particles as core–shell structures with a refractive index typical of extracellular vesicles (shell RI = 1.48, core RI = 1.38) [14], [15], [19]. No significant differences in size distribution were detected; all liposome populations exhibited a consistent median diameter of approximately 72 nm and a log-normal distribution (Figure 2E, figure S#?). To further assess resolution, 150-fluorophore liposomes and blanks were combined and analysed following compensation using single-stain controls. tSNE analysis provided clear separation of the distinct populations, and bivariate fluorescence plots confirmed discrete clustering of liposome populations labeled with different fluorophores (Figure 2F–G). Complementary imaging flow cytometry (ImageStream X MkII) was performed for comparison, enabling visualization of individual fluorescent liposomes. Here, only the preparations containing 150 fluorophores were readily distinguishable from unstained liposomes, with Atto595-labeled liposomes demonstrating the most robust signal, highlighting the enhanced sensitivity of the CytoFLEX Nano (Figure S2A–C). Collectively, these findings demonstrate that the CytoFLEX Nano possesses sufficient sensitivity to resolve small lipid nanoparticles differing in fluorescence intensity and for accurate multiplex detection.

**Figure 2.**
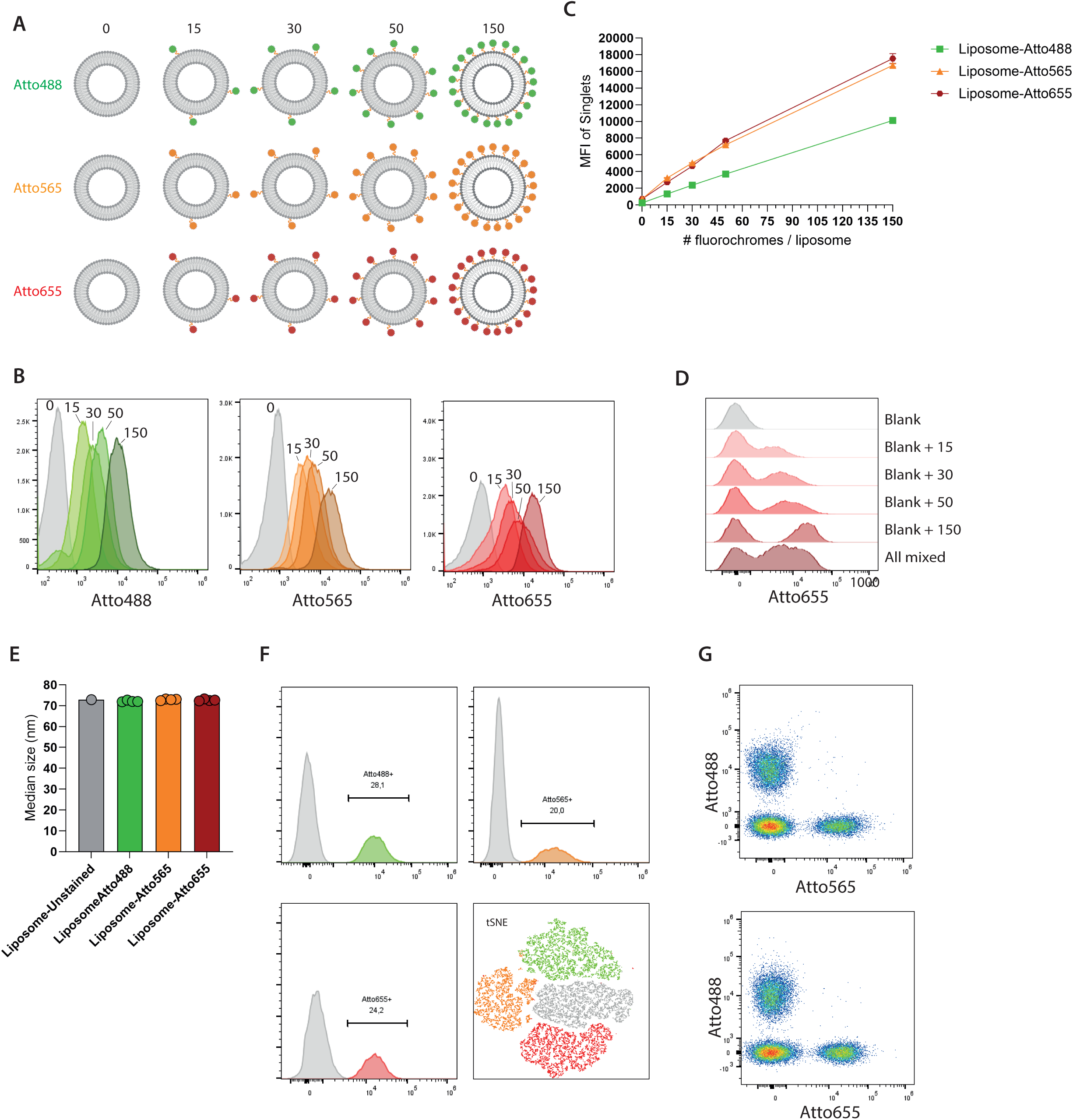
Liposomes with differing numbers of fluorochromes. A) Overview of the liposomal samples. B) Overlayed histograms of individual liposome samples showing separation of the peak fluorescence. C) Median fluorescence intensity (MFI) quantification of all liposome samples. D) Mix of blank liposomes and fluorescent liposomes showing the separation of positive and negative peaks. E) Median size of liposomes derived from scatter calibration with FCMpass software. F) Mix of blank with 150-fluorochrome liposomes of each color, and separation of these on histograms and tSNE plot. G) Separation of fluorescent liposome population on bivariate plots.

### Background matters

Given the high sensitivity of the CytoFLEX Nano, we next sought to systematically evaluate background particles and potential sources contributing to elevated event rates in flow cytometry samples. Interestingly, PBS was found to contain numerous small and occasional larger events, with particle content varying according to vendor and lot. Notably, changes in PBS lot over a one-year period coincided with marked increases in larger event counts within the established inclusion gate (Figure 3A). This observation underscored the importance of filtration; using Whatman Anotop 20 nm syringe filters for PBS effectively reduced particle counts by over 80% in the inclusion gate (Figure 3B), compared to unfiltered PBS or sheath fluid. Although filtration of the dilution buffer mitigates background, it may also be necessary to filter biological samples prior to analysis, as the manufacturer (Beckman Coulter) recommends excluding particles above 1 μm in diameter. However, filtration with a 0.45 μm PES syringe filter resulted in elevated counts of larger events in both PBS and peritoneal lavage samples (Figure 3C-D). Application of a chemical isolation reagent (Total Exosome Isolation

**Figure 3.**
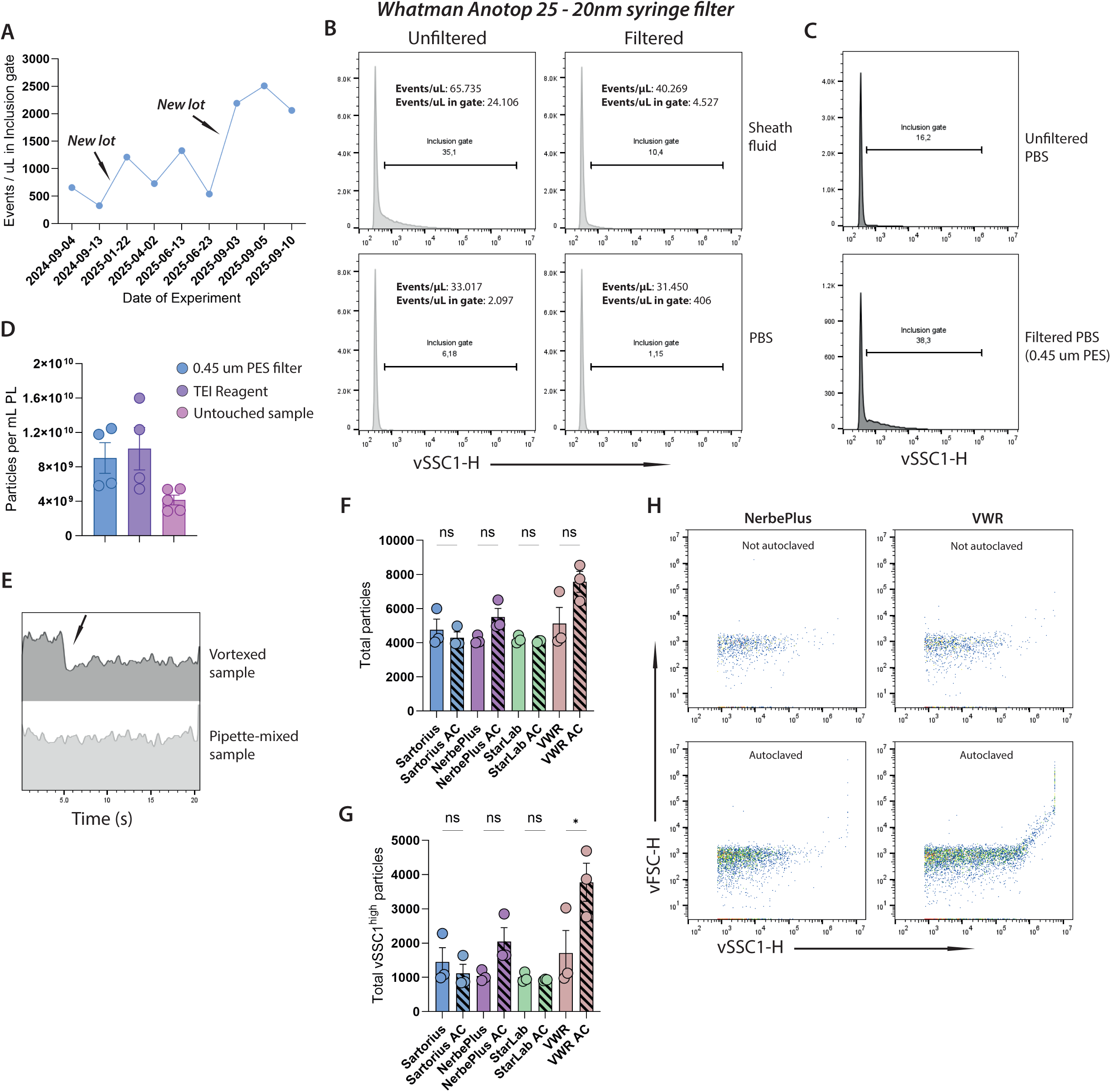
Background particles from materials and buffers. A) Particle concentration in PBS from the inclusion gate from different lot numbers of PBS and different experiments spanning a one-year period. B) Histograms of sheath fluid or PBS which are either unfiltered or filtered through a 20 nm Whatman Anotop 25 syringe filter. C) Histograms of unfiltered and 0.45 µm PES syringe filtered PBS D) Bar graph showing the particle concentration peritoneal lavage samples that were filtered through 0.45 µm PES filter, isolated with total exosome isolation reagent, or untouched. E) Histogram time-plot showing the effect of vortex-mixed samples compared to pipette-mixed sample. F) Bar graph showing total particle counts from non-autoclaved vs autoclaved tubes from four different brands. G) Bar graph of total vSSC1^high^ particles between non-autoclaved and autoclaved tubes from four different brands. H) Bivariate plots of representative non-autoclaved and autoclaved samples from brands that had increased particles after autoclavation.

Reagent) similarly increased particle numbers relative to untreated controls (Figure 3D). Additional methodological variables were found to influence background event rates. Vortex-mixed samples exhibited pronounced initial surges of background events relative to pipette-mixed counterparts (Figure 3E), and the type of analysis tube used marginally affected overall background signal (Figure S3A). Furthermore, some brands of pipette tips inflated total particle numbers when autoclaved, while other brands did not (Figure 3F-H). The pipette tips affected by autoclavation also produced a higher number of particles with higher scattering intensity (vSSC1^high^) which bleed into the inclusion gate (Figure 3G, Figure S3BC). We also noticed that autoclaved 1.5 mL Eppendorf tubes contained a lot of particles compared to 5 mL Falcon tubes, which could be eliminated by filtering through 20 nm filter (Figure S3D). In summary, multiple factors, including buffer composition, sample filtration, reagent use, sample handling, and tube selection, can elevate background particle events. Experimental strategies to minimize background should be prioritized, and appropriate controls are essential to reliably account for background contributions.

### Qualitative comparison of membrane dyes

Selecting the most suitable membrane dye for EV analysis or EV–cell uptake studies is often challenging, with limited consideration given to dye quality or optimal concentration. The CytoFLEX Nano represents a useful platform for evaluating and optimizing the choice and application of membrane dyes. To this end, we compared four representative dyes: a CTB-based dye (ExoBrite CTB which is cholera toxin b labelling GM1 gangliosides) and three lipophilic dyes (DiO, ExoBrite True EV Membrane Stain, and MemGlow640). To assess aggregate formation, peritoneal lavage samples were stained in technical triplicates, including detergent and PBS + stain controls. DiO labelling resulted in substantial variability between the triplicates, and similar particle counts in the PBS + stain control indicated considerable aggregate or micellar formation at the concentration used (Figure 4A). Both ExoBrite True and MemGlow640 yielded comparable particle counts between samples and showed minimal aggregate formation in PBS. However, ExoBrite CTB stained substantially fewer particles, possibly reflecting subtype-specific labeling (Figure 4A). To maximize EV detection, a dual-labeling strategy with ExoBrite True and ExoBrite CTB was adopted. Dyes were stored at 4–8°C as recommended, and over a one-year period, quality controls from the same stocks revealed a gradual increase in the number of stained particles within the EV gate (Figure 4B). However, when analyzing paired samples of PBS and PBS + stain controls from two different time points, we could see that the increase in stained particles coincided with an increase in background particles from the PBS (Figure 4B). By looking at the EV gate from the different PBS samples, no stain was present in PBS with the low background, whereas more stained particles were detected in PBS with high background (Figure 4C). This indicates an interaction between the lipophilic dye and particles in PBS which potentially increases the background staining and further underscores the importance of including staining controls for sample analysis.

**Figure 4.**
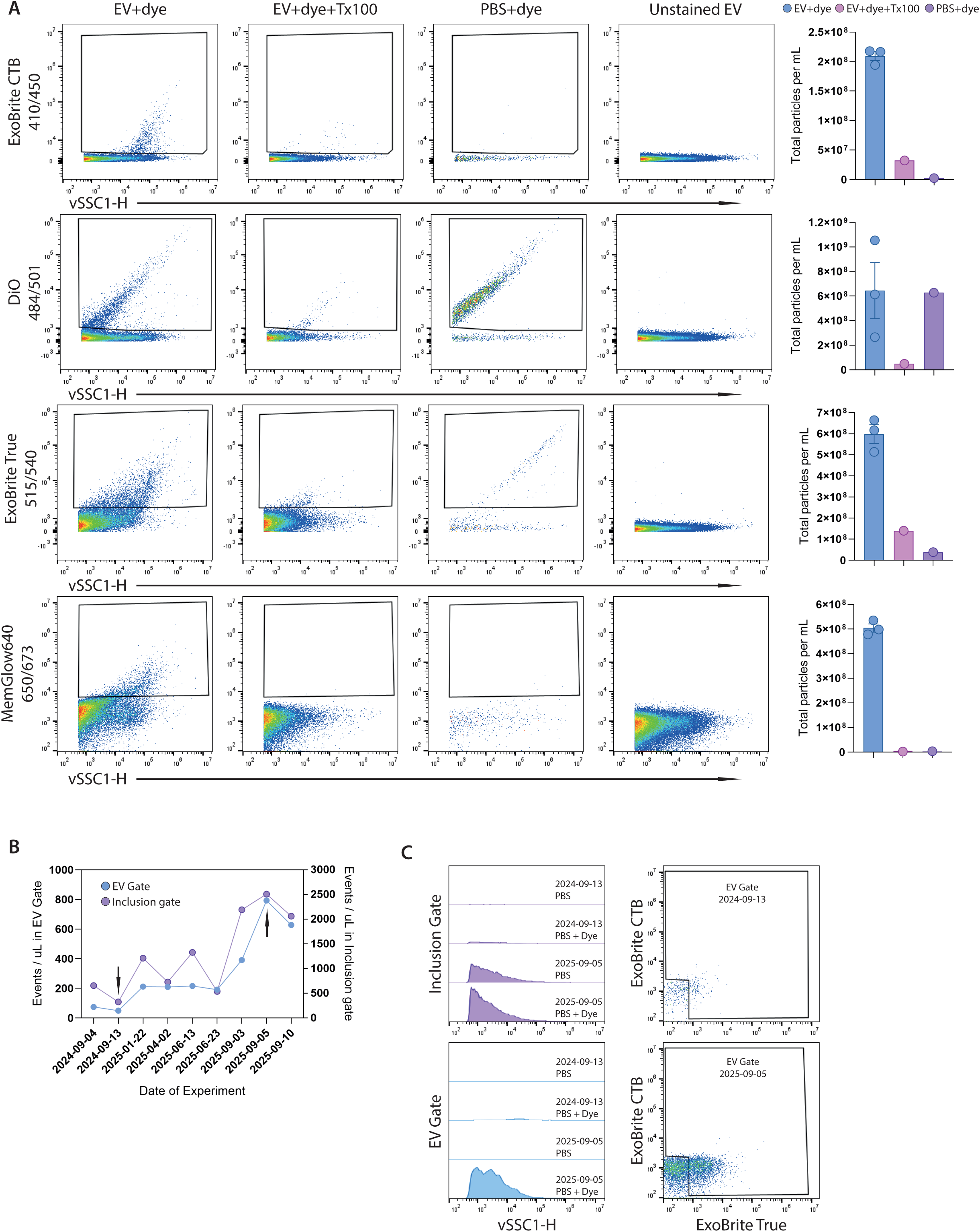
Qualitative assessment of membrane dyes for EV staining. A) Bivariate plots of the different dyes after staining samples from mouse peritoneal cavity fluid. Detergent controls, PBS + stain control, and unstained controls are all displayed as well as a quantitative graph showing the total number of particles per mL. B) XY graph showing particle concentration in PBS (right Y axis) and PBS + stain (left Y axis) over the course of different experiments. Arrows indicate the lowest and highest counts, respectively. C) Offset histograms and bivariate plots of PBS and PBS + stain from two different experiments performed one year apart. The ExoBrite True reagent was from the same lot in both experiments.

### Characterization of single extracellular vesicles from peritoneal fluid and cell culture medium

After characterizing the sources for increased background and potential solutions, we next investigated the capability of the instrument in analyzing biological samples. As input, we used fluid from peritoneal lavage and set out to compare EVs from unprocessed or ultracentrifuged samples in technical quadruplicates (Figure 5A). Due to the sensitivity of the instrument and the low sample input, we tested how manual handling and dilution affects event rates between the samples and found an almost identical event rate (ungated) between the 8 samples tested. This suggests a high stability and reproducibility of the instrument (Figure 5B). Next, we sought to distinguish EVs from other particles and stained samples using the two different membrane stains, ExoBrite CTB and ExoBrite True. We set the inclusion gate based on the PBS control to exclude the background peak, then gated on single particles, and ultimately on EVs including both single and double positive for the two membrane stains (Figure 5C). When comparing the EV frequency between ultracentrifuged samples and samples with no isolation we found a ∼14-fold relative EV enrichment (Figure 5D). As expected, we also found a decrease in particle concentration (∼8-fold) in the ultracentrifuged samples compared to the no isolation samples (Figure 5E). Zooming in on the concentration of EVs we saw a less dramatic reduction of roughly ∼2-fold, translating to an EV recovery of ∼55 % (Figure 5F). Surprisingly, we saw that ExoBrite CTB-positive EVs were not precipitated by ultracentrifugation which could explain the low EV recovery compared to the no isolation samples (Figure 5G). Collectively, these findings highlight both the robustness of the instrument for EV detection and the risk of methodological biases introduced by a common EV isolation workflow; in our case, a failure to recover ExoBrite CTB-positive EVs in the ultracentrifuged fraction.

**Figure 5.**
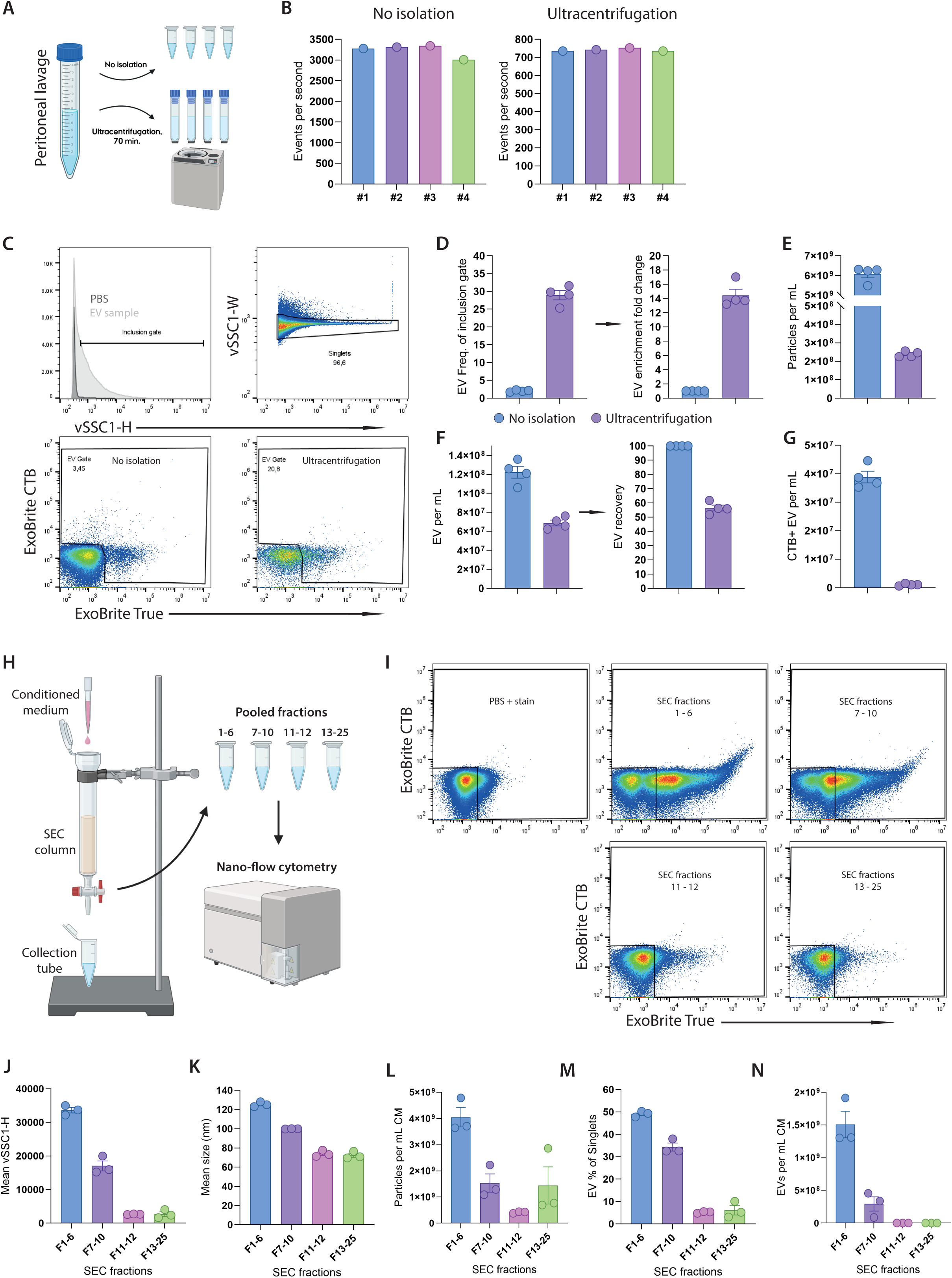
Interrogation of EVs from biological samples. A) Schematic overview of isolation procedures for peritoneal fluid samples. B) Bar graph showing event rate (events per second) for untouched and ultracentrifuged samples. C) Gating strategy of biological samples, with an inclusion gate being placed according to the PBS control, then singlets are gated out based on vSSC1 width and height, and then an EV gate is set based on the two membrane dyes ExoBrite CTB and ExoBrite True. Events positive for either of the two or both are regarded as EVs. D) Bar graphs showing the frequency (%) of EVs in inclusion gate between untouched and ultracentrifuged samples. The fold change was then calculated and plottet as relative EV enrichment. E) Bar graph of particle concentration between untouched and ultracentrifuged samples. F) Bar graph presenting EV concentration for untouched and ultracentrifuged samples. The percentage difference is shown as relative EV recovery. G) Bar graph showing the concentration of ExoBrite CTB+ EVs in untouched and ultracentrifuged samples. H) Schematic overview of SEC separation of conditioned medium. I) Bivariate plots to show how the EV gate is set between fractions. J) Mean scatter (vSSC1-H) intensity of the pooled fractions. K) Scatter calibrated data for all pooled fractions plotted as diaemeter (nm). L) Bar graph of particle concentration of each pooled fraction. M) Percentage of EVs of the singlet (inclusion) gate. N) Concentration of EVs in all pooled fractions.

We also analysed EVs from cell culture supernatants isolated using SEC. As done in previous studies, 25 fractions of 0.5 mL per sample were collected and pooled into four groups: fractions 1-6, 7-10, 11-12, and 13-25 with fractions 7-10 typically representing EV-enriched fractions [20], [21] (Figure 5H). As for the peritoneal lavage samples, pooled fractions and controls were stained with the same ExoBrite reagents and analysed individually (Figure 5I). We used the same initial gating as with the peritoneal samples (inclusion gate, singlets) and set the EV gate according to the PBS + stain control and applied it to all fractions (Figure 5I). Despite a clear separation between the positive and negative populations as seen on the ExoBrite plot in F1-6 and F7-10, we decided to keep the same EV gate across all samples to avoid potential overestimation of stained particles (Figure 5I). In SEC, the largest particles are eluted in the first fractions, reflected in decreasing side scatter intensity from the first to the last fractions (Figure 5J).

To estimate the size of these EVs we performed scatter calibration as described for the liposomes above. In the pooled EV-fractions, we found that F1-6 had a mean diameter of ∼125 nm, F7-10 ∼100 nm, and both F11-12 and F13-25 had a mean diameter of ∼70 nm (Figure 5K). The concentration of single particles was highest in F1-6 and the lowest in F11-12 with F7-10 and F13-25 being intermediate and more similar, indicating a higher concentration of large compared to small particles (Figure 5L). We then analysed the frequency of EVs in all fractions and found that F1-6 had ∼50 % EVs, F7-10 ∼35 % EVs, and the last two fractions were below 5 % (Figure 5M). Finally, to estimate the concentration of EVs we subtracted the background events and found that F1-6 contained >3-fold more EVs than F7-10 and that F11-12 and F13-25 contain negligible amounts of any EVs (Figure 5N). Once again, we illustrated the stability and sensitivity of the instrument, while demonstrating that the particle size, concentration, and EV frequency all declined across fractions, with EVs largely confined to the earliest pools as expected.

### Sizing and enumeration of extracellular vesicles

Estimating particle concentration with CytoFLEX Nano is based on dividing the total number of particles by measured flow compared to other platforms where calibration beads are used. This is facilitated by the highly precise piston pump in the CytoFLEX Nano that allows for accurate volumetric estimation resulting in reproducible estimations of concentration. We initially compared the concentration estimates of peritoneal lavage determined using the NTA and the CytoFLEX Nano and found that NTA reported ∼4.6-fold higher particles per mL compared to the CytoFLEX Nano using statistics from the ‘inclusion gate’, which corresponds to the analytical level of the NTA samples (Figure 6A). With each date representing a different experiment, we tracked whether the percentage of EVs in peritoneal lavage was stable over time (Figure 6B). Surprisingly, the EV percentage of total particles was quite reproducible around the 3 % (Figure 6B). Similar reproducibility was reflected in the number of CD9^+^ EVs per mouse (3-4 x 10^7^) (Figure 6C).

**Figure 6.**
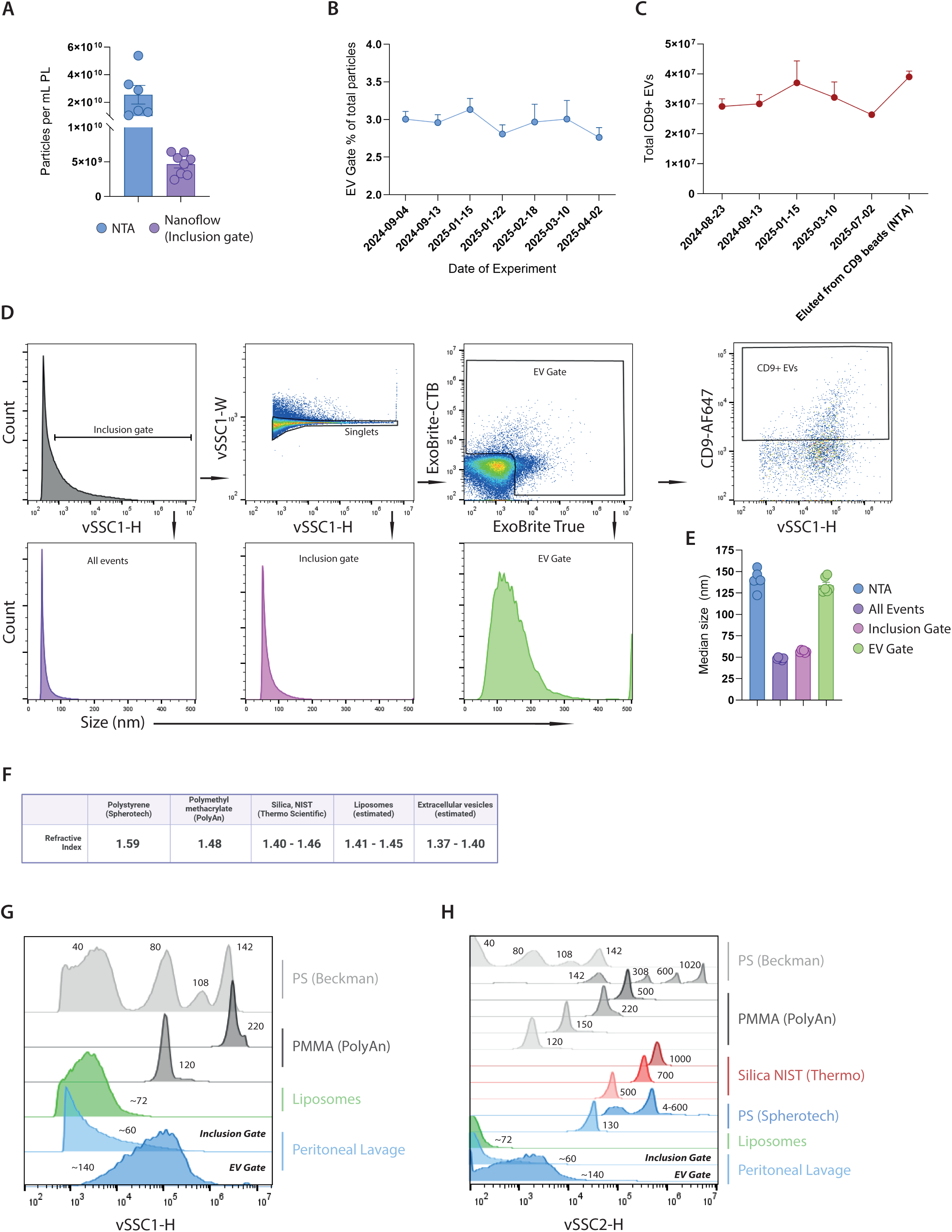
Reproducibility and sizing. A) Bar graph showing particle concentration in PL measured by NTA and Nanoflow using the inclusion gate as input. B) XY graph showing the percentage of peritoneal EVs from healthy mice tracked over several experiments. C) XY graph of CD9+ EV concentration in PL measured over time. D) Gating strategy for the samples and a histogram of scatter calibrated data within each of the gates, except the CD9 gate. E) Bar graph showing the median size of particles analysed with NTA compared to median size of particles within each of the gates. F) Table of refractive index of the different particles used in this study. G) Offset histograms showing the scatter distribution (vSSC1-H) of the smaller particles and EVs. H) Offset histograms showing the scatter distribution of all particles on the vSSC2-H.

Next we estimated the size of particles using FCMPASS in three different gates from the CytoFLEX Nano: All events (no gating), inclusion gate (singlets, %background), and the EV gate based on the two membrane stains ExoBrite CTB and the lipophilic ExoBrite True EV Membrane Stain combined (Figure 6D). For the ‘all events’ gate and inclusion gate, the sizes of the events were ∼50 and ∼55 nm, whereas the size of events in the EV gate was ∼140 nm (Figure 6E). We compared these sizes to particle sizes obtained during NTA analysis of the same type of samples and saw that the NTA reported ∼145 nm sized particles although no sub-gating or enrichment was done prior to the analysis (Figure 6E).

Lastly, we compared how various sizes of beads made of different materials and thus with different RI looks on the CytoFLEX Nano. We analysed NanoVIS polystyrene beads from Beckman Coulter, Spherotech polystyrene beads, PMMA beads from PolyAn, Silica beads from Thermo Scientific, and compared those to our in-house made liposomes and peritoneal lavage samples (Figure 6F-H). We checked scatter distribution in the sensitive vSSC1 for the smallest particles, showing a clear effect of the RI of the different materials (Figure 6G). For example, 80 nm PS beads have similar peak intensity to 120 nm PMMA beads and ∼140 nm PL EVs, and the peak of 40 nm PS beads was higher than that of ∼72 nm liposomes (Figure 6G). The same was observed for the ‘large particle’ detector vSSC2, where the scatter distribution of the different samples reflects the RI (Figure 6H). Taken together, the CytoFLEX Nano demonstrates great reproducibility of concentrations and has the resolution to separate small particle populations based on their RI and scatter intensities.

## Discussion

This study demonstrates the power, reproducibility, and capability of the CytoFLEX Nano to resolve small nanoparticles and EVs. We confirm that the instrument can easily separate PS beads down to 40 nm in size based on scatter intensity, which greatly exceeds the 80 – 100 nm limit of detection reported for conventional flow cytometers [12], [13]. As has been shown in several other publications, RI correction is essential when estimating EV size from scatter intensity [22]. A study illustrated that 80 nm PS beads had similar side scatter intensities as ∼158 nm EVs on the instrument used which indicates that small EVs (>200nm) are close to the detection limit of conventional flow cytometers [23]. When comparing 80 nm PS beads to peritoneal EVs using the CytoFLEX Nano, we also saw a similar result with the 80 nm PS bead peak coinciding with our EV peak at ∼140 nm (Figure 6G). This could indicate that if the limit of detection of a flow cytometer corresponds to 100 nm PS beads, the vast majority of small EVs below 200 nm in size would not even be detected by scatter intensity and would, thus, require fluorescence labelling (Figure 6G). In terms of fluorescence, we show that the instrument can detect as few as 15 fluorochromes per liposome while we could only reliably detect 150 fluorochromes per liposome using an imaging flow cytometer optimized for small particle detection. However, a recent study reported a detection limit of 10 Phycoerythrin (PE) fluorophores on two different conventional flow cytometers [24]. PE is one of the brightest fluorochromes available with a calculated brightness index of ∼1.660.000 (calculated as Extinction Coefficient x Quantum Yield) versus 72.000 (Atto488), 108.000 (Atto565) and 37.500 (Atto655), translating into PE being 23.1, 15.4, and 44.3-fold as bright, respectively. As the CytoFLEX Nano could identify 15 Atto fluorochromes per liposome it could theoretically detect down to 1 PE fluorochrome. In future studies, it would be of great interest to determine the limit of detection of the CytoFLEX Nano for different fluorochromes.

As conventional flow cytometers are primarily designed for cell analysis, hardware components are optimized to maximize signal-to-noise ratio while preserving high-throughput measurement capacity for cell-based assays. Here, interference between buffer and reagent-derived nanoparticles would not affect analysis due to the threshold and gain set for size of cells. For nanoparticle interrogation the background becomes more prominent since it contains particulates of similar size and scattering intensities as the sample nanoparticles/EVs. We found that the increased sensitivity of the CytoFLEX Nano truly brings the background detection to light: Everything from vortexing the samples before analysis, the type and lot number of dilution buffers, what filters, tubes, and reagents are used – all could introduce significant amounts of background particles that fall within the inclusion gate. The same applies for membrane dyes and antibodies which can contribute with background signals appearing/giving signals in the different subgates, such as EV gates. For example, we show that DiO produced a lot of fluorescent particles at the applied concentration with an almost linear dependency between scatter and fluorescence intensity in both stained EV samples and in PBS + stain, indicating the presence of lipid-dye micelles (Figure 4A). Although similar tendencies for aggregates were observed for ExoBrite True, it was much less pronounced, indicating that a more appropriate concentration or a dye that is less prone to emit fluorescence when in micellar formations could be considered. When plotting fluorescence against vSSC1-H the shape of stained samples of ExoBrite True and MemGlow640 differed from those of DiO, not having the distinct linear correlation between scatter and fluorescence, thus looking more like expected heterogeneous EVs. Furthermore, we observed that an increase in background particles in PBS correlated with higher numbers of ExoBrite True positive events, suggesting that particles present in PBS may associate with the lipophilic dye and subsequently bleed into the EV gate, artificially increasing the apparent EV counts (Figure 5B). Hence, rigorous staining optimization and controls are necessary to avoid/minimize or at least reliably correct for background or reagent-induced particles.

Additionally, we assessed how the instrument handles two different isolation procedures of natural biological samples. For peritoneal fluid, we could run untouched samples (no isolation) which allowed us to assess EV recovery and EV enrichment of ultracentrifuged samples. EV recovery after ultracentrifugation was approximately 55% compared to untouched samples, however, none of the single-positive ExoBrite CTB events were detected in the ultracentrifuged samples, and this loss was the primary contributor to the low recovery observed after ultracentrifugation. Although GM1 gangliosides (the ligand for ExoBrite CTB) have been implicated in EV biogenesis [25], the absence of ExoBrite CTB-positive events in our ultracentrifuged samples may be explained either by the possibility that these events are not EVs (since they were also negative for the lipophilic dye) or by their low density, which could make them more difficult to precipitate by ultracentrifugation.

To further validate the instrument, we analysed samples fractionated by SEC. We pooled fractions that were likely to contain EVs and fractions likely to be absent of EVs and analysed them on the CytoFLEX Nano. Here, we could verify that the largest particles were present in the first fractions and smallest in the last fractions, and that EVs were present in fractions 1-10 and absent in 11-25. In some cases, fractions 1-6 are treated as ‘void’ and thus not analysed, whereas fractions 7-10 are considered EVs [26]. In our case, pooled fractions 1-6 contain the most particles and most EVs closely followed by pooled fractions 7-10. For future studies, it would be beneficial to further dissect the SEC fractions and include assessment of EV-associated proteins and lipoproteins for each fraction to rule out possible contamination of lipoproteins.

For physical characterization of EVs NTA has been the most widely used for more than a decade [27]. NTA is a high-throughput, easy to use instrument which provides a relatively objective estimation of nanoparticle size and concentration; however, it also tends to have a strong bias towards larger particles since they scatter more light [28]. In our study, we found that the NTA tends to estimate nanoparticle size in peritoneal fluid to be around ∼145 nm, whereas the CytoFLEX Nano only showed sizes between 50 – 55 nm, and it was only when we used the EV gate for size input that the size was more similar to NTA. Despite being biased towards the larger particles, we still saw a drastic difference of particle concentration by NTA compared to the CytoFLEX Nano. In contrast, another study showed that NTA underestimated EV/particle numbers compared to another nano flow cytometer [28]. Generally, after EV isolation, all particles detected by NTA (or other quantification method) are referred to as EVs – neglecting contribution from non-EV particles and buffer/reagent-derived particles. We show here that common EV enrichment methods like ultracentrifugation and SEC still contain a lot of non-EV particles. We saw that only ∼30 % and 35 – 50 % particles that fall within the EV gate, respectively, indicating that more than half of the sample is non-EVs. In the non-isolated peritoneal fluid samples, only 3 – 5 % are within the EV gate, further underscoring that EVs in most cases are rare events in biological fluids [11]. These frequencies do not take various lipoprotein particles, which can also be stained by lipophilic dyes (Simonsen, 2017) into account. This clearly highlights the importance of high-resolution phenotyping of individual particles to properly discriminate between true EVs and other nanoparticles present in a sample. The CytoFLEX Nano, or nano flow cytometry in general, could in theory also be used to discriminate between large lipoprotein particles and EVs assuming proper antibodies against the different apolipoproteins have been validated for flow cytometric purposes.

## Future perspectives

Nano flow cytometry is a high-throughput, single-particle, multiparametric technique that enables phenotyping and quantification of EVs, surpassing traditional bulk characterization methods like NTA or EM in multi-parameter capability. Flow cytometry has revolutionized phenotyping of single cells, and we believe that the continuous development and advancement of dedicated nanoparticle flow cytometers will revolutionize phenotyping of single EVs and other biological nanoparticles. Once the first instruments have been stress-tested and procedures thoroughly validated, development of nanoparticle sorters and spectral modules for nanoparticle flow cytometers would further increase the resolution, versatility, and robustness of EV research as we have seen for the cell-based counterparts.

## Concluding remarks

In this study, we demonstrate that the CytoFLEX Nano can accurately resolve small nanoparticles of different materials based on side scatter and fluorescence measurements. We also present a robust analysis of EVs from two distinct sources, isolated using two different methods, and compare their scattering properties (size) with those of calibration beads of different materials. The CytoFLEX Nano is a powerful instrument that can greatly enhance high-throughput EV analysis with unprecedented resolution.

## Acknowledgements

The authors wish to acknowledge the FACS Core Facility at Aarhus University. The authors like to thank Thomas Wittenborn and Bettina Winther Grumsen for technical assistance.

---

**Figure S1.**
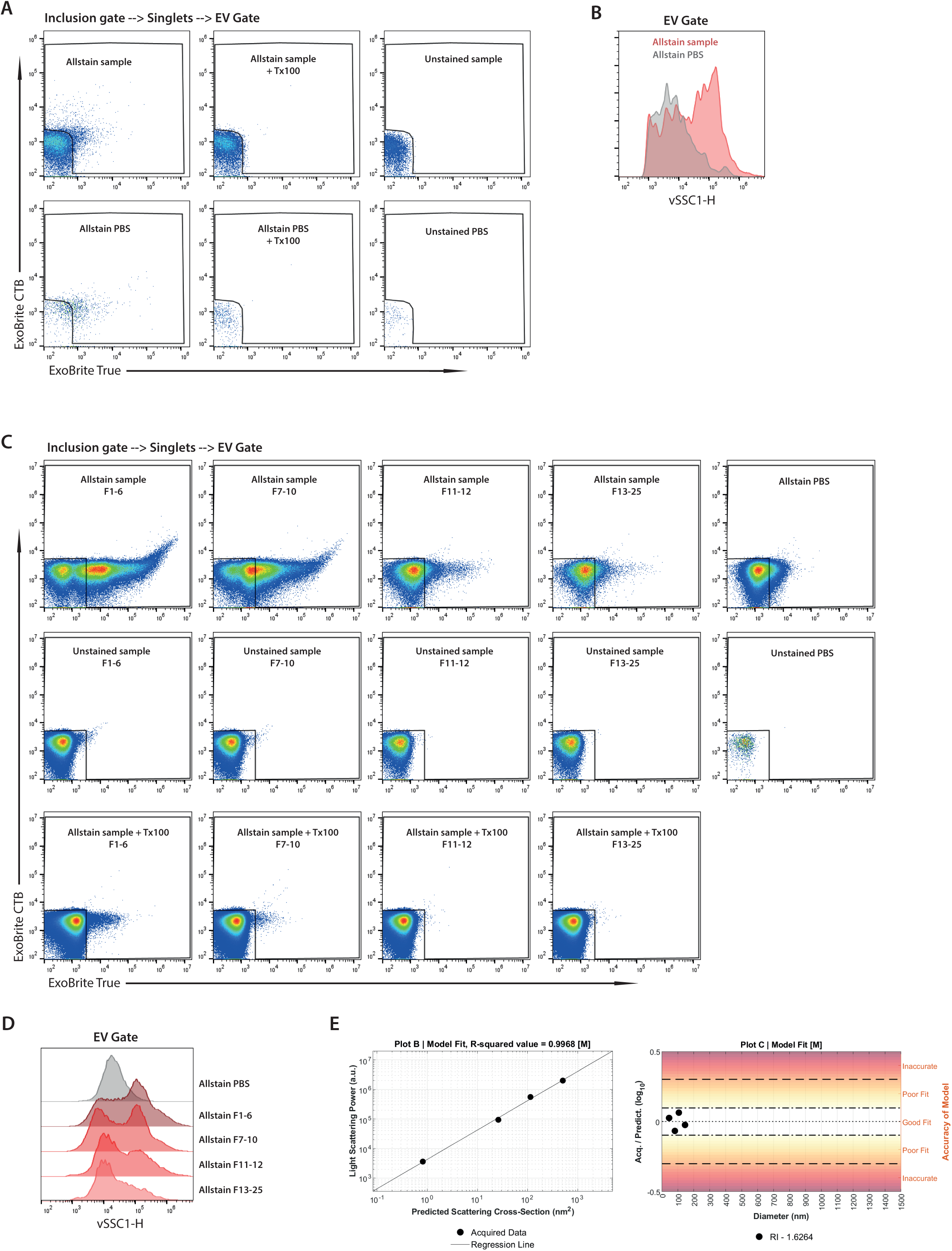
Controls for biological samples. A) Bivariate plots of EV gate from peritoneal fluid samples showing representative stained sample and PBS, detergent treated stained sample and PBS, and unstained sample and PBS. B) Histogram showing the difference in scatter profile on the vSSC1-H of stained PBS and stained sample from peritoneal fluid. C) Bivariate plots of EV gate from conditioned medium and SEC fractionated samples showing representative stained sample and PBS, detergent treated stained sample and PBS, and unstained sample and PBS for all pooled fractions. D) Histogram showing the difference in scatter profile on the vSSC1-H of stained PBS and stained sample from conditioned medium. E) Plots showing the accuracy of the prediction model from FCMPASS using NanoVIS Low beads as a model test.

**Figure S2.**
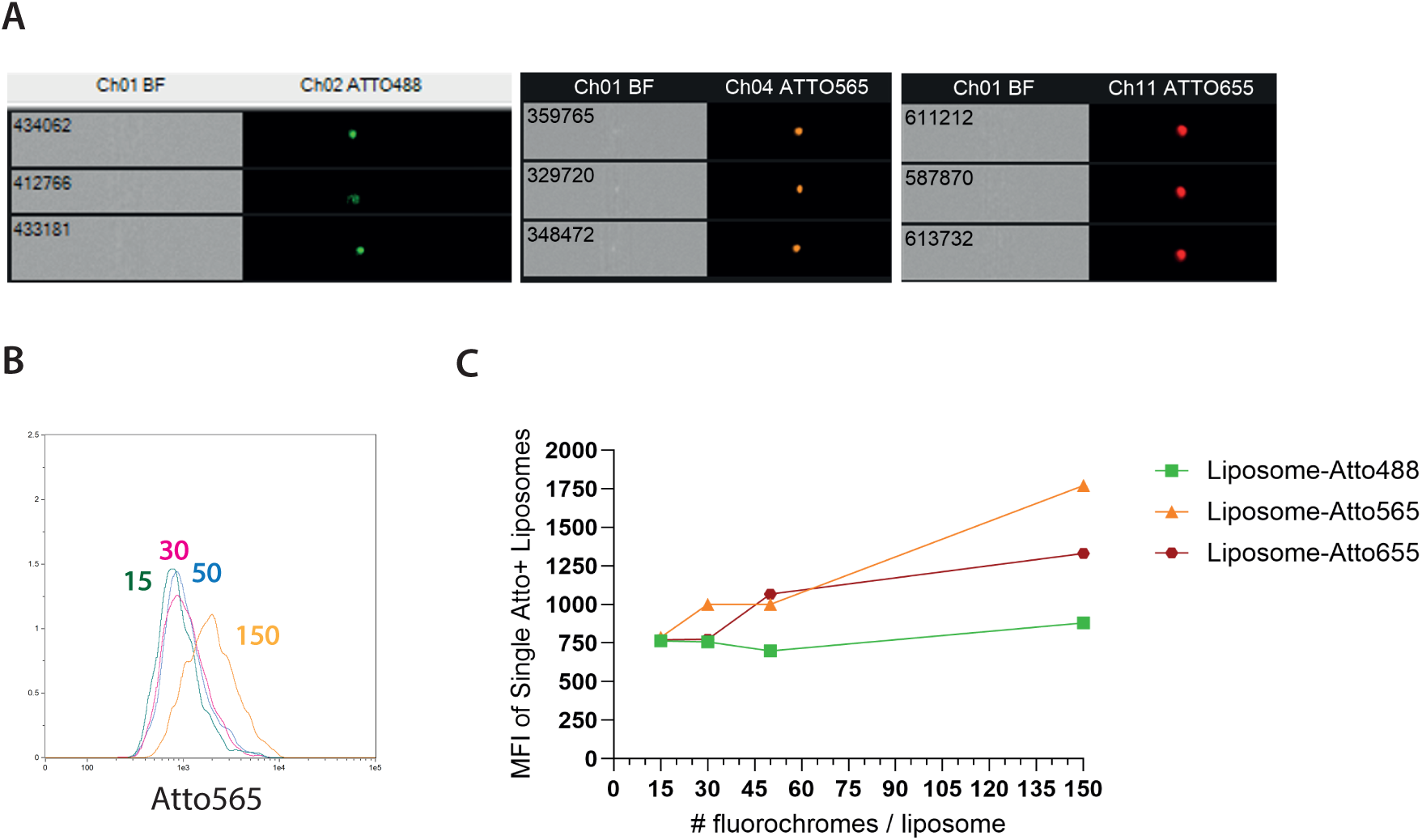
Imaging flow cytometry. A) Single fluorescent events detected of liposomes with the different fluorochromes. B) Histograms of liposomes with different numbers of fluorochromes. C) XY graph showing the MFI of the different liposome samples.

**Figure S3.**
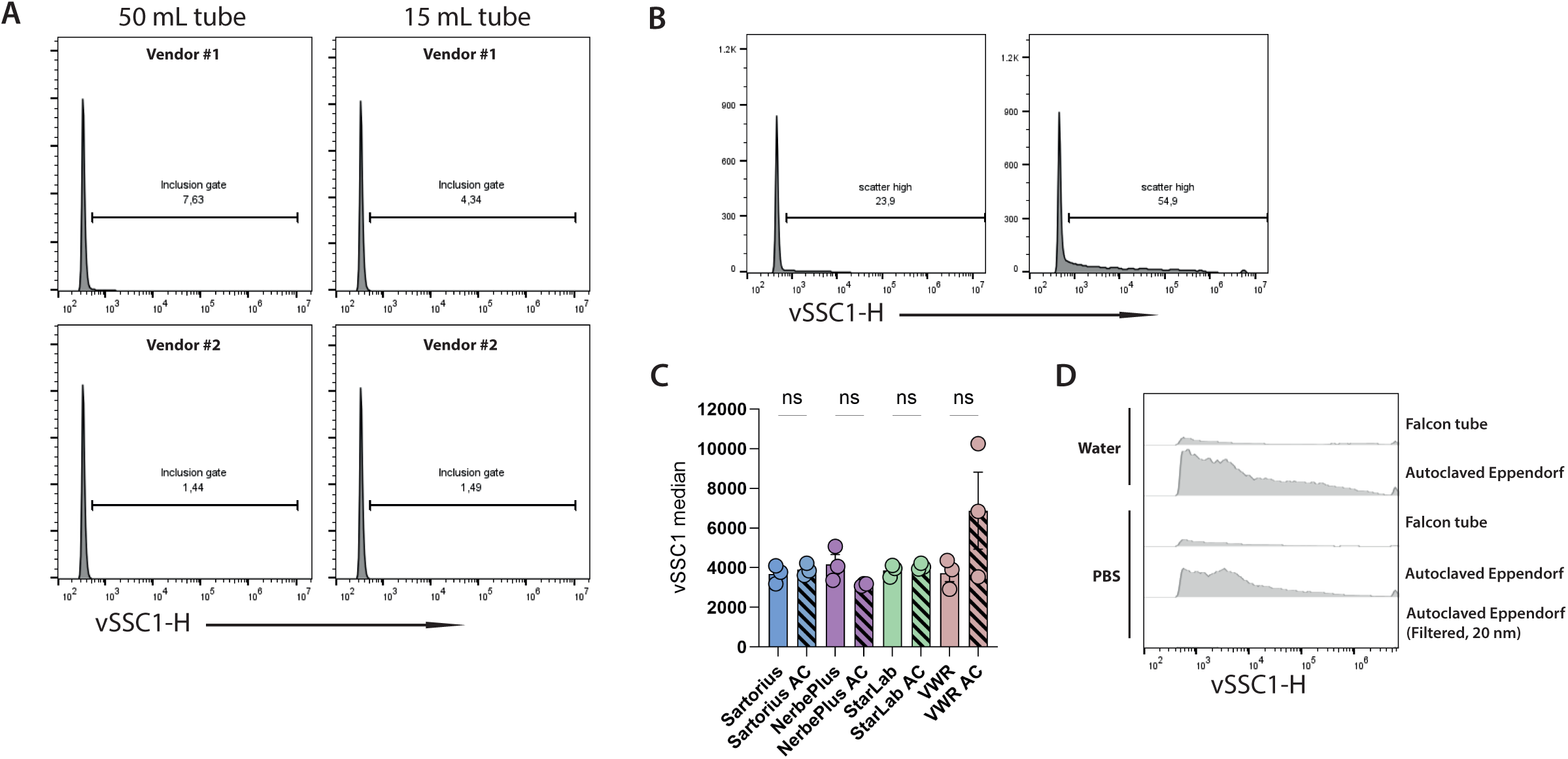
Background matters. A) Number of inclusion gate particles from tubes bought from vendor 1 compared to vendor 2. B) Histograms showing inclusion gate (vSSC1^high^ particles). C) Bar graph showing the median scatter intensity of the vSSC1^high^ particles. D) Offset histogram showing the differences between a 5 mL Falcon tube (used for all analyses) versus 1.5 mL autoclaved Eppendorf tube.

